# A reference-free strategy for circulating tumor DNA detection from whole-genome sequencing data

**DOI:** 10.1101/2025.07.23.666290

**Authors:** Carmen Oroperv, Amanda Frydendahl, Tenna Vesterman Henriksen, Giovanni Santacatterina, Alice Antonello, Nicola Calonaci, Mads Heilskov Rasmussen, Giulio Caravagna, IMPROVE-consortia, Claus Lindbjerg Andersen, Søren Besenbacher

**Author notes:** Membership of the IMPROVE-consortia is provided in the Acknowledgments.

## Abstract

Circulating tumor DNA (ctDNA) is emerging as a promising biomarker for postoperative monitoring of cancer patients. Precise estimation of circulating tumor fraction is crucial for evaluating treatment effects and timely detection of disease recurrence. All current ctDNA detection methods that utilize whole-genome sequencing (WGS) data rely on the reference genome alignment of sequencing reads and often apply separate tools for detecting different variant types. However, various bioinformatic analysis confounders and the application of external variant calling tools could be avoided by analyzing k-mers from unaligned sequencing reads. While k-mer-based methods have successfully been applied for somatic variant validation and detection, the potential of k-mer-based ctDNA detection is unexplored. We have developed a tumor-informed reference-free ctDNA detection tool called *ctDNAmer* that detects tumor-specific somatic variation directly from unaligned sequencing data by identifying k-mers unique to the tumor DNA. ctDNAmer detects variant information across the genome by comparing the primary tumor and germline WGS data and accounts for sample-specific germline variability and technical noise in the same framework. We tested the utility of ctDNAmer for tumor fraction estimation on postoperative plasma cfDNA WGS data (mean sequencing depth ~28x) from 90 stage III colorectal cancer patients with three years of follow-up. The tumor fraction (TF) estimates agreed with the available clinical information and ctDNA was detected in 77% (17/22) of recurring patients with a median lead time of 8 months compared to radiological imaging. We further validated ctDNAmer’s tumor fraction estimates based on a comparison with the mean cfDNA allele frequencies of somatic clonal SNVs identified from aligned primary tumor sequencing data. The TF estimates showed a strong Pearson correlation of 0.897 with the mean allele frequencies and improved ctDNA detection results across samples with an AUC of 0.79 compared to 0.75 if the mean allele frequency of clonal mutations is used.

## Introduction

Circulating tumor DNA (ctDNA) is poised to transform cancer care, emerging as a promising biomarker for post-treatment surveillance. ctDNA fragments, originating from the primary tumor, circulate in the bloodstream of cancer patients and can provide a direct view of the cancer genome and tumor dynamics. Unlike invasive tissue biopsies, ctDNA can be sampled from blood plasma [1], and its short half-life enables real-time tumor burden monitoring [2]. Furthermore, ctDNA analysis can capture the temporal and spatial heterogeneity of the tumor, offering a more comprehensive molecular profile than single-site tissue biopsies [3]. These features make ctDNA an advantageous alternative to standard surveillance methods, such as radiological imaging and blood-based metabolic cancer biomarkers. Several studies have demonstrated that ctDNA testing could improve recurrence monitoring [4–6] and may be beneficial for guiding treatment decisions [7–9].

Detection of ctDNA in low tumor burden settings, such as minimal residual disease (MRD), presents significant challenges and requires highly sensitive methods capable of identifying small quantities of tumor-derived DNA amidst the vastly larger background of cell-free DNA (cfDNA) originating predominantly from healthy hematopoietic cells [10]. In tumor-informed approaches, where primary tumor tissue is available, the circulating tumor fraction (fraction of ctDNA fragments out of all cfDNA fragments) can be measured by identifying tumor-specific mutations in cfDNA. Methods for targeted mutation detection can be divided into two main categories: PCR-based techniques, such as digital PCR [11], and next-generation sequencing (NGS)-based approaches, including Duplex sequencing [12] and CAPP-Seq [13]. These methods have demonstrated high sensitivity, achieving detection of allele fractions as low as 0.01% [12–14].

While targeted approaches are highly sensitive, untargeted whole-genome sequencing (WGS)-based analysis offers distinct advantages for ctDNA detection. Targeted approaches usually use much higher sequencing depth than untargeted whole-genome (WGS) data, and will thus have better sensitivity to detect a specific mutation, but recent studies show that WGS data can compensate for this by including a much larger number of variants in the analysis [15]. Furthermore, WGS enables tumor-specific mutation detection to be extended from single nucleotide variants (SNVs) to other variant types, such as copy number alterations [16], or carry out multi-feature analyses that incorporate additional ctDNA characteristics, such as fragment length profiles [17].

In the tumor-informed setting, ctDNA detection from WGS involves two main steps: (1) identifying tumor-specific somatic variants from the primary tumor sequencing data [18] and (2) searching for these patient-specific mutations in cfDNA [19,20]. The conventional approach for somatic variation detection relies on aligning sequencing reads to the reference genome, followed by applying variant calling tools to identify alternative alleles by comparing aligned reads to the reference sequence [21]. However, short-read alignment is a complex process that requires heuristic algorithms and can be confounded by biological and technical factors. Biological factors that can confound alignment include genetic variability not represented in the reference genome [22–24], segmental duplications, and repetitive sequences [25]. Technical issues, such as the incompleteness of the reference genome [26], DNA damage that has occurred during sample preparation, and sequencing errors [27], further complicate alignment. These confounding effects may lead to unmapped reads, resulting in data loss, or ambiguous or wrongly mapped reads, which can result in mismatched bases mistakenly identified as somatic mutations in downstream analyses [28–30].

Variant calling can be further complicated by low tumor purity of the tumor sample [25,31], or if the primary tumor is sequenced from formalin-fixed paraffin-embedded (FFPE) tissue. FFPE samples are prone to higher levels of DNA damage from the fixation process compared to fresh-frozen (FRFR) tissue [32,33]. In addition, despite algorithmic advancements, different variant-calling methods often produce discordant results. Several studies have demonstrated that the precision and recall of called variants strongly depend on the combination of the read mapping algorithm and variant calling tool used [25,34–37]. This is especially true for other mutation types besides SNVs and variants present in a small fraction of cells [28,38,39].

All the above-mentioned confounding factors associated with the bioinformatic analysis of WGS data can affect tumor fraction estimation, and no standardized workflow has yet been established. However, the use of raw whole-genome sequencing reads for ctDNA detection remains largely unexplored, even though reference-free and/or alignment-free k-mer-based approaches have been applied for variant detection and validation in other settings [40–48].

In this study, we present a reference-free approach for ctDNA detection, *ctDNAmer*, that detects tumor-specific somatic variation directly from unaligned sequencing data by identifying k-mers unique to the tumor sample. These k-mers are then used to detect ctDNA within raw cfDNA sequencing reads. ctDNAmer leverages genome-wide information from the primary tumor and is not limited to SNVs. The method identifies tumor-specific k-mers by comparing primary tumor and germline WGS data, accounts for patient-specific germline variability and technical noise in the same framework, and applies probabilistic modeling to estimate the circulating tumor fraction. We validate ctDNAmer by estimating tumor fractions of stage III colorectal cancer (CRC) patients’ postoperative plasma cfDNA samples. The estimated tumor fractions are compared against radiological follow-up results and mean cfDNA allele frequencies of clonal SNVs identified from aligned tumor sequencing reads, and the results clearly demonstrate that it is possible to derive good-quality tumor fraction estimates using a reference-free approach.

## Results

### 1. Reference-free tumor-specific somatic variation and ctDNA detection

Tumor-specific somatic variants define the uniqueness of the tumor genome when compared to the genomes of the patient’s healthy cells. The core idea of ctDNAmer is to capture this uniqueness with k-mers (DNA sequences of length k) extracted directly from raw sequencing reads, bypassing the need for read alignment to the reference genome and somatic variant calling (Figure 1a and 1b). A fraction of these unique tumor k-mers should then be detectable in cfDNA samples if the cancer cells shed tumor DNA into the bloodstream.

**Figure 1.**
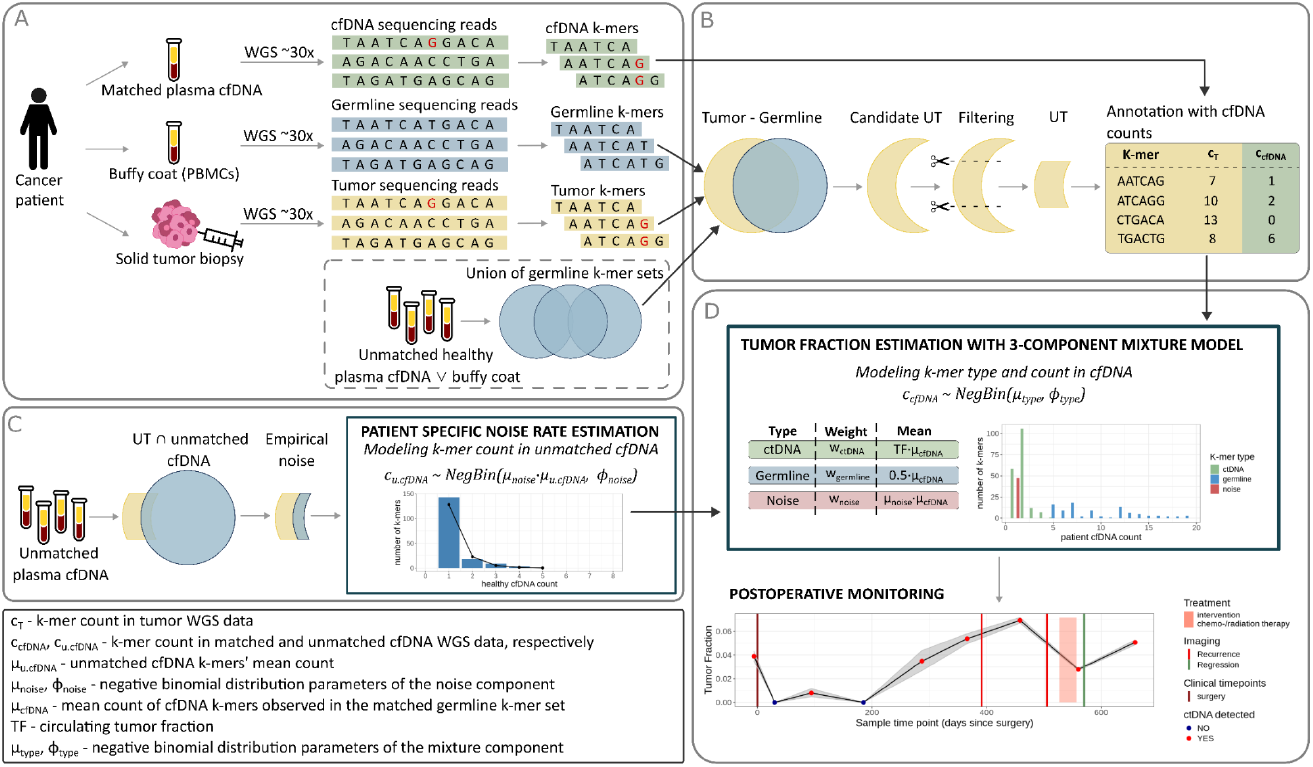
Overview of the ctDNAmer workflow. **a)** K-mer counting from three data sources and the germline union creation across healthy cfDNA and buffy coat samples (dashed box). The nucleotide bases marked with red in the tumor and cfDNA sequences denote tumor-specific somatic variants; **b)** Identification of unique tumor (UT) k-mers by primary tumor and germline sets comparison, candidate UT set filtering and UT k-mers annotation with cfDNA counts; **c)** Empirical noise rate estimation from unmatched cfDNA samples; **d)** Tumor fraction estimation based on the UT k-mer counts in the matched cfDNA. Colored circles indicate k-mer sets. Overlaid circles indicate set operations between sets. UT: unique tumor. “Unmatched healthy plasma cfDNA” refers to cfDNA from healthy individuals, “unmatched cfDNA” refers to cfDNA from any individual other than the target patient (can include both healthy cfDNA and patient cfDNA).

An overview of ctDNAmer can be seen in Figure 1. ctDNAmer first counts overlapping k-mers (k = 51) from the raw sequencing reads of the tumor, matched germline (DNA from peripheral blood mononuclear cells (PBMCs)), and cfDNA to create input k-mer sets for ctDNA detection. An initial candidate set of k-mers unique to the tumor (UT k-mers) is created by subtracting germline k-mers from the tumor k-mer set. This candidate UT set is further filtered and annotated with the cfDNA count data. ctDNAmer then models the observed count of these UT k-mers in the cfDNA as a function of the circulating tumor fraction (TF), assuming that the counts and relative abundance of UT k-mers in cfDNA increase proportionally with the tumor fraction.

Besides tumor-specific variants, two alternative explanations can account for k-mers appearing unique to the tumor and detectable in the cfDNA. First, a k-mer may have been present in both the tumor and germline genomes, but remained undetected in the germline data, and thus was misclassified as a UT k-mer. Second, a k-mer could be erroneously labeled as UT due to technical noise. Such k-mers capturing technical noise are present in neither the tumor nor germline genomes but result from tumor DNA sequencing errors (possibly caused by DNA damage).

To accommodate these two alternative explanations of UT k-mers, ctDNAmer models UT k-mers’ cfDNA count with a three-component mixture model: 1) ctDNA, 2) germline, and 3) technical background noise (Figure 1d). Distinct count distributions of the three components enable ctDNAmer to distinguish tumor-specific k-mers from germline and noise k-mers. If the UT k-mer is a genuine tumor-specific k-mer, its count in the cfDNA is expected to be proportional to the tumor fraction. In contrast, the counts of k-mers explained by the missed germline component or the technical noise component are independent of the tumor fraction. Germline k-mers are expected to have relatively high counts close to the sample coverage, or half of the coverage if they capture heterozygous variants, reflecting their origin from the more abundant healthy cfDNA fragments. Conversely, technical noise k-mers, occurring randomly on DNA fragments and sequencing reads, are expected to have a lower mean count compared to germline k-mers.

To inform the mixture model of the expected noise level, ctDNAmer first estimates the parameters of the sample-specific noise distribution based on empirical data (Figure 1c), which are then used as fixed input parameters in the mixture model (see next section, Figure 1d). The possibility of a missed germline variant is handled by estimating a sample-specific rate of missed germline k-mers in a pre-operative cfDNA sample and then using this fixed rate in the analysis of all subsequent samples.

### 2. Sample-specific background noise estimation

After germline subtraction and k-mer filtering, which includes removing k-mers with low or high counts and/or GC content to increase the signal-to-noise ratio (see Methods section 3.1 for details), we expect most k-mers in a UT set to represent somatic tumor variation. However, given the stochastic nature of technical noise in sequencing data, we need to account for potential residual noise during TF estimation to prevent inflation of TF estimates and false positive ctDNA detections.

We assume that each UT set has a specific technical noise level that is mainly determined by the tumor sample damage levels and sequencing error rate. If stochastic noise consists of k-mers randomly generated by the tumor DNA technical alterations, a fraction of these k-mers should be observed by chance in any unmatched cfDNA sample. Hence, ctDNAmer utilizes cfDNA samples from other individuals to estimate the set-specific background noise rate. These could be cfDNA samples from healthy donors with no ctDNA, or cfDNA samples from other patients. Unmatched patient cfDNAs are acceptable controls, as UT k-mer sets are patient-specific. Applying several cfDNA samples for the noise rate estimation ensures reliable parameter estimates that reflect the average rate of UT k-mers observed in cfDNA if no ctDNA is present. Supplementary Figure 1 shows that using 15 healthy cfDNA samples in the CRC cohort resulted in a strong correlation of mean k-mer counts between the healthy samples and matched ctDNA-negative postoperative samples (Pearson correlation coefficient of 0.82 for k-mers with a count of one), which is indicative of accurate noise rate estimation. Nevertheless, the optimal number of samples for noise rate estimation may vary according to target sequencing depth and error rate variance among the unmatched samples.

The empirical noise data set is created by intersecting cfDNA k-mer sets from other individuals with the UT set. The cfDNA counts of the intersections k-mers’ are then used to estimate the mean and variance of the background noise distribution (see Methods section 5 for details). The noise distributions are modeled using a negative binomial distribution to account for possible overdispersion in the count data, and the distribution mean is scaled by the mean k-mer count of the corresponding cfDNA samples to ensure adjustment for varying sequencing coverages.

### 3. Tumor fraction estimation for postoperative monitoring of stage III colorectal cancer patients

To demonstrate ctDNAmer’s utility for TF estimation, we applied it to plasma cfDNA samples from 90 stage III colorectal cancer (CRC) patients. Primary tumor FRFR tissue and matched germline WGS data were available for each patient. The plasma cfDNA samples were collected during a three-year follow-up period after resection of the primary tumor, with one preoperative sample also available from each patient. Radiological imaging detected disease recurrence for 22 patients during the three-year follow-up period.

We identified candidate UT k-mers from the primary tumor data by subtracting the matched germline k-mer sets and a panel of germline k-mers created from a larger set of samples, as leveraging germline information from multiple individuals has been shown to reduce false positives in somatic variation detection [28]. The union of germline k-mer sets was created from the germline WGS data of 45 CRC patients (included in the analyzed cohort) and cfDNA samples of 30 healthy donors (see Methods section 3 for details). This union provided a more comprehensive representation of the germline genome sequences and removed additional k-mers from all candidate UT sets (on average, this filtering removed 69% of the tumor k-mers that remained after the matched set was subtracted).

Noise rates of the UT sets were estimated based on empirical data created from k-mer sets of 15 healthy donor cfDNA samples (not included in the germline union). To test the assumption that unmatched cfDNA samples and matched ctDNA-negative samples have similar numbers of observed UT k-mers, which can be accounted for by the background noise distribution, we compared the mean number of UT k-mers in the donors’ cfDNA to the mean number of k-mers in ctDNA-negative cfDNA of the non-recurring patient (Supplementary Figure 1). The mean numbers showed strong correlations for k-mers observed once or twice in the cfDNA samples (Pearson correlation coefficients of 0.82 and 0.69, respectively), consistent with the expectation of TF-independent background noise. Further, there was substantial variability in the mean number of UT k-mers observed in donors’ cfDNA across the sample cohort (Supplementary Figure 1, range of 3 to 200 for k-mers observed once). This variability can be explained by differing error rates and damage levels of the underlying tumor samples and emphasizes the need for set-specific noise rate estimation. Supplementary Figure 2 shows the mean and variance of the estimated noise distributions, assuming an equal cfDNA mean count of 30.The parameter estimates reflect the variability in the noise rates and help to ensure that differing noise levels will be accounted for during TF estimation.

Estimates of the cfDNA samples’ TFs were obtained from the three-component mixture model, representing true tumor, germline and noise k-mers’ distributions. As the mixture component weights were assumed to remain constant, the preoperative cfDNA samples were used to estimate the germline component weight and distribution parameters. These estimates were subsequently applied as fixed input parameters for TF estimation in postoperative cfDNA samples. Similarly, the preoperative sample’s tumor component weight estimate was set as the mean of the prior distribution in all subsequent cfDNA samples (see Methods section 6 for details).

The ctDNA status ground truth was established based on radiological imaging data. To assess the ability of ctDNAmer to detect the presence of ctDNA in a sample, we first looked at a subset of samples that we could confidently label as either ctDNA-positive or ctDNA-negative. We labeled all preoperative cfDNA samples and the recurring patients’ postoperative samples collected before treatment following the recurrence as ctDNA-positive. Postoperative samples from non-recurring patients, collected after the treatment (either surgery or surgery followed by adjuvant chemotherapy, which was administered for 81 of the 90 patients), were labeled as ctDNA-negative. In total, the ctDNA status was known for 718 cfDNA samples, comprising 207 positive and 511 negative samples.

We compared the ground truth to the ctDNA status determined based on the ctDNAmer’s TF estimates. The ctDNA detection cutoff was set at the TF value that maximized Youden’s J statistic (maximum Youden’s J: 0.51 (Figure 2a); see Methods section 7 for details). Figure 2b illustrates the TF values of all ctDNA-positive and -negative samples along with the detection cutoff value of 0.0006, where a clear difference in TF estimates of the ctDNA-positive and -negative sample groups can be observed (ctDNA-positive group TF mean 0.0275, ctDNA-negative group TF mean 0.0006; Mann-Whitney-Wilcoxon test p-value < 1.05*10^−34^).

**Figure 2.**
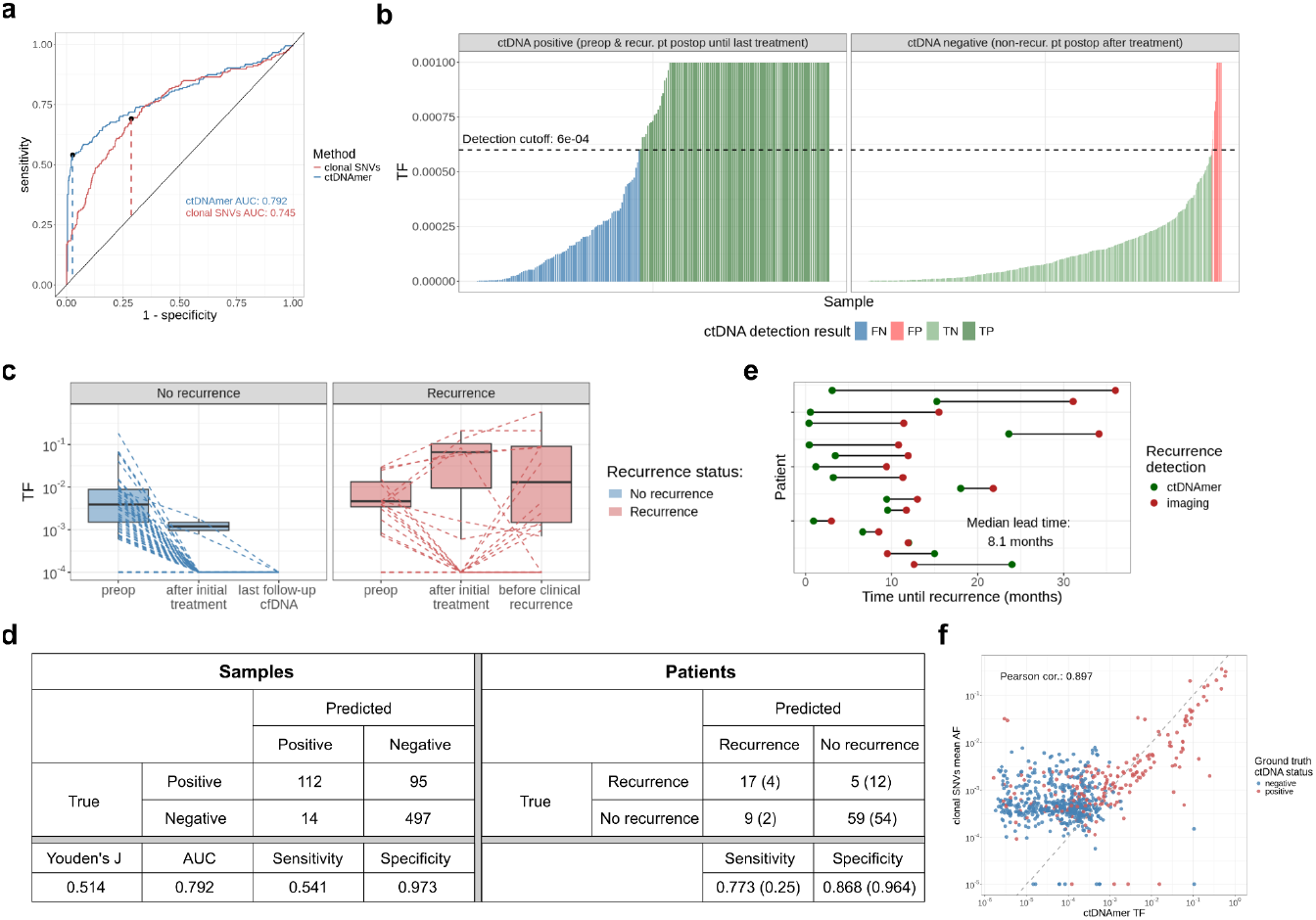
ctDNA detection results in postoperative cfDNA samples of 90 stage III colorectal cancer patients. **a)** ROC curve. The dashed lines indicate maximum Youden’s J statistic values; **b)** Estimated tumor fractions of ground truth ctDNA-positive and -negative samples. Samples are colored based on their ctDNAmer detection result. The dashed black line indicates the detection cutoff chosen based on maximizing Youden’s J. FN: False negative; FP: False positive; TN: True negative; TP: True positive; **c)** Comparison of TF estimates of recurring and non-recurring patients at different categorical time points; dashed lines indicate TF estimates of each individual across the time points; y-axis is on a logarithmic scale, zero values were excluded from the boxplot calculation and set to 10^−4^ for the dashed lines; **d)** Confusion matrices of detection results on the sample and patient level. The values outside the parentheses in the patient matrix represent serial analysis that includes all samples collected during the follow-up period. The values inside the parentheses in the patient matrix represent landmark analysis where only one sample, the first sample collected after the surgery and before the start of the adjuvant chemotherapy, was analysed from each patient (72 out of the 90 patients had the landmark sample available); **e)** Comparison of recurrence detection times between ctDNAmer and radiological imaging. The ctDNAmer’s recurrence detection time point was chosen as the first postoperative ctDNA-positive cfDNA sample, even if adjuvant chemotherapy was still ongoing; **f)** Comparison of ctDNAmer’s TF estimates and mean allele frequencies of clonal SNVs. Only samples with known ground truth status determined based on clinical data were included in the correlation calculation and figure f, axes are on a logartihmic scale, clonal SNV mean allele frequencies of zero were changed to 10^−5^.

Figure 2c gives an overview of the TF estimates of recurring and non-recurring patients at different time points. As expected, the preoperative estimates are similar across the two patient groups. However, the TF estimates of the first cfDNA sample obtained after the treatment completion (surgery or surgery + adjuvant chemotherapy) show variation between the two groups. While estimates of non-recurring patients are close to zero, indicating no ctDNA presence, a fraction of recurring patients show elevated TFs. As expected, recurring patients show elevated TF estimates also at the time point of the last cfDNA sample obtained before recurrence was confirmed with radiological imaging. The non-recurring patients’ TF estimates stay close to zero across the follow-up period, as illustrated by the low estimates of the last cfDNA samples collected during the follow-up period.

Using ctDNAmer and the chosen detection cutoff, ctDNA was detected in 57 (63%) preoperative cfDNA samples. Figure 2d presents the ctDNA detection results in postoperative cfDNA samples on the sample level (individual ctDNA-positive and -negative samples, as defined above) and patient level (detection in any postoperative cfDNA sample for the recurring patients, and no detection across samples in non-recurring patients). At the sample level, we observed a high specificity of 0.973. However, specificity was decreased to 0.868 at the patient level when analysing all serial samples, as one or two false positive detections were observed among postoperative samples of several patients. These false positives may have been caused by specific cfDNA samples’ elevated error rates, not observed during the background noise estimation in the empirical data. The k-mer-based approach showed a moderate sample-level sensitivity of 0.54. Sensitivity improved to 0.77 at the patient level for serial samples analysis, as recurrence detection was possible even if not all postoperative samples were ctDNA-positive. The landmark sample – the first cfDNA sample obtained after the surgery and before the start of adjuvant chemotherapy – is clinically highly relevant. Emerging as a marker for residual disease, it can provide valuable insight for risk assessment and treatment decisions following the tumor resection. The classification performance for this sample across patients is presented in parentheses in the patient-level confusion matrix in Figure 2d. The landmark sample was available for 72 out of the 90 patients. ctDNAmer showed high detection specificity of 0.96 for the landmark samples. However, the detection sensitivity dropped to 0.25. This may partly be explained by the chosen detection cutoff, as Youden’s J was maximized based on all serial samples. Further, the varying landmark sample timepoints can have an effect on detection sensitivity due to the presence of trauma-induced cfDNA. It has been shown that in CRC patients, the total cfDNA level is increased postoperatively on average threefold up to four weeks after surgery [49]. Trauma-induced cfDNA increase can make it more difficult to detect the minute amounts of ctDNA present after the tumor resection. While all 72 patients’ landmark samples were obtained at least 7 days after the surgery, only five patients had the landmark sample timepoint later than 28 days after surgery, which could partly explain the lower detection sensitivity.

Based on ctDNAmer’s TF estimates and the selected detection cutoff, we were able to detect ctDNA in 17 (77%) recurring patients. When defining recurrence detection time as the first ctDNA-positive postoperative cfDNA sample, ctDNAmer achieved a median lead time of 8.1 months compared to radiological imaging (Figure 2e), detecting recurrence earlier for 14 out of the 17 patients. These results suggest that if ctDNA is detected, ctDNAmer may provide an opportunity for earlier clinical action, such as an earlier timepoint for radiological imaging, coupled with potential earlier intervention for the recurrence. Supplementary Figure 3 displays TF estimates of all cfDNA samples from recurring patients. In general, estimates decrease between the preoperative and the first postoperative cfDNA sample, indicating that surgery removed the vast majority of the cancer cells that shed DNA. Among patients where ctDNAmer was able to detect recurrence, TF estimates show a continuous increase before the time point when recurrence was radiologically confirmed (Supplementary Figure 3). After an intervention following recurrence, the TF estimates decrease as expected (Supplementary Figure 3). Either persistent low TF estimates and ctDNA-negative status or a new increase in TF estimates and ctDNA-positive classification can be observed, which may be indicative of the treatment effect (Supplementary Figure 3).

Lastly, we investigated how ctDNAmer’s performance is affected by the k-mer length. In addition to k-mer length of 51, we tested ctDNAmer on the same CRC patient cohort with k-mer lengths of 31, 41 and 61. ROC curves of ctDNA detection results are shown in Supplementary Figure 4a. The best AUC was achieved with the k-mer length of 51 and performance decreased both when decreasing and increasing the k-mer length. Nevertheless, the decrease in the AUC values was small and ROC curve characteristics did not change remarkably. The lowest AUC of 0.767 was observed for k-mer length of 31, demonstrating the robustness of ctDNAmer’s performance across k-mer lengths.

### 4. ctDNAmer’s TF estimates comparison with clonal SNVs allele frequencies

To validate ctDNAmer’s TF estimates and ctDNA detection performance, we analysed the same CRC patient cohort cfDNA samples with a customary alignment-based approach for tumor-informed ctDNA detection. We identified clonal tumor-specific SNVs from aligned sequencing data and tracked the allele frequency of these variants in the postoperative cfDNA samples (see Methods section 8 for details). Assuming that ctDNAmer correctly detects ctDNA k-mers and that the counts and relative abundance of these k-mers depend on the TF, we expected ctDNAmer’s TF estimates to correlate with the mean cfDNA allele frequency of clonal SNVs.

We set the ctDNA detection cutoff for the mean allele frequencies in the same way as for ctDNAmer’s TF estimates, choosing the value that maximizes Youden’s J. Comparing ctDNA-detection on the sample level, ctDNAmer achieved better detection results compared to clonal SNVs (Figure 2a, ctDNAmer AUC: 0.792, clonal SNVs AUC: 0.745). The recurring patients’ cfDNA mean allele frequencies, along with the ctDNAmer’s TF estimates, can be seen in Supplementary Figure 3. The two methods estimates align across most of the recurring patients’ cfDNA samples, indicating agreement in disease dynamics for most recurrences and treatment effects. TF estimates for all cfDNA samples where ctDNA status was known based on clinical data (see Methods section 7) can be seen in Figure 2f, where a strong correlation between the two methods’ estimates is shown (Pearson correlation of 0.89). The agreement of the two orthogonal measures further proves the validity of ctDNAmer for TF estimation.

### 5. ctDNAmer’s TF estimates comparison with ddPCR and Signatera allele frequencies

While ctDNAmer’s TF estimates strongly correlate with clonal SNVs allele frequencies, both approaches are based on the same WGS data set. To further validate ctDNAmer’s TF estimation performance, we compared the estimates with allele frequencies determined by droplet digital PCR (ddPCR) [50] and Signatera [51]. These frequency estimates were obtained from paired aliquots of the same cfDNA plasma samples from a subset of patients. The comparison results can be seen in Supplementary Figure 5. Paired estimates from ddPCR and ctDNAmer were available for 270 cfDNA samples. ctDNAmer showed slightly improved ctDNA detection with an AUC of 0.782 compared to ddPCR performance with an AUC of 0.773 (Supplementary Figure 5a). Paired estimates from Signatera and ctDNAmer were available for 569 cfDNA samples, and Signatera showed superior performance with an AUC of 0.864 compared to ctDNAmer’s AUC of 0.777 (Supplementary Figure 5c). ctDNAmer’s TF estimates showed strong correlation with both ddPCR and Signatera allele frequencies with Pearson correlation coefficients of 0.931 and 0.905, respectively (Supplementary Figures 5b and 5d). Both the ctDNA detection performance comparison and the TF estimates’ correlation with ddPCR and Signatera further validate the utility of ctDNAmer for TF estimation based on unique tumor k-mer counts in WGS cfDNA data.

### 6. Tumor fraction estimation based on FFPE tumor-derived UT k-mers

We also evaluated ctDNAmer’s performance on an FFPE tumor-tissue cohort of 24 stage III CRC patients. With a k-mer length of 51 and using an FFPE-specific detection cutoff set by maximizing Youden’s J statistic, ctDNAmer achieved improved detection sensitivity in the FFPE cohort compared to FRFR, both at the sample level (FFPE: 0.75, FRFR: 0.54) and patient level (FFPE: 1.00, FRFR: 0.77). However, specificity was notably lower in the FFPE cohort, particularly on the patient level (FFPE: 0.32; FRFR: 0.87). Notably, ctDNAmer’s performance for the FFPE tumor sample patient cohort improved remarkably with shorter k-mer lengths (Supplementary Figure 4b). The best detection performance was observed with a k-mer length of 31, which was the shortest tested length. With k equal to 31, ctDNAmer achieved an AUC of 0.782 for the FFPE cohort (Supplementary Figures 6c and 6f), which is comparable to the detection performance of the FRFR patient cohort with a k-mer length of 51 (AUC of 0.792). The difference in the optimal k-mer length for these two cohorts can be explained by increased fragmentation of DNA extracted from FFPE samples compared to FRFR samples [52].

The FFPE cohort ctDNA detection results with a k-mer length of 31 can be seen in Supplementary Figure 6. Given the increased damage levels typically observed in FFPE samples compared to FRFR tissue [32,33], we expected to observe higher noise rates in UT sets derived from FFPE samples. Supplementary Figure 6a and 6b do not confirm this assumption but indicate equal noise rates in the two cohorts. FFPE cohort mean noise rates (mean of 0.0067 across samples) are comparable to the FRFR UT sets’ noise rates (mean of 0.0064 across samples). This can be due to successful candidate UT sets’ filtering, where k-mers capturing technical noise were removed. The TF estimates at the three clinically relevant categorical time points can be observed in Supplementary Figure 6e. Similarly to FRFR patients, non-recurring FFPE patients have low estimated TFs at the first postoperative time point and at the last available cfDNA sample time point. Recurring patients show reduced TFs after the surgery and increased TFs before clinical recurrence, as expected. Compared with the FRFR cohort results with k-mer length of 51, improved sensitivity of 0.61 at the sample level and of 1.00 at the patient level was achieved for the FFPE cohort with the k-mer length of 31 (Supplementary Figure 6f). However, detection specificity was reduced both at the sample level (0.82) and patient level (0.58). Similarly, compared to the FRFR cohort, increased sensitivity and reduced specificity was also observed for the landmark sample analysis of the FFPE cohort (Supplementary Figure 6f). Although there were only five recurring patients in the FFPE cohort, a lead time of 5.8 months, reduced by two months compared to the FRFR cohort, was achieved (Supplementary Figure 6g).

## Discussion

In this study, we demonstrated that tumor-informed ctDNA detection can be performed directly from unaligned sequencing data without relying on reference genome alignment or external variant calling tools. ctDNAmer identifies k-mers unique to the tumor genome by subtracting known germline sequences and uses these k-mers to estimate the circulating tumor fraction of cfDNA. By bypassing read alignment, ctDNAmer prevents data loss from unaligned or misaligned reads and data pre-processing biases. Instead, k-mers of interest are chosen from the candidate set based on the sequence composition and input data distribution. Our analysis of 90 colorectal cancer patients’ data showed that tumor fraction estimates obtained with ctDNAmer strongly correlate with the mean allele frequency of somatic variants identified from aligned sequencing data. However, additional validation on diverse WGS datasets, including different sequencing depths and cancer types, is needed to further establish the generalizability of the method.

A key advantage of ctDNAmer is its ability to detect all tumor-specific k-mers regardless of the number of differences to the corresponding germline sequence. This enables simultaneous detection of SNVs, indels, structural variants, and complex variants where multiple mutations, possibly of different types, have occurred in close proximity. Capturing all variant types within a single framework may improve detection sensitivity compared to approaches that rely solely on predefined variant types.

Compared to radiological imaging, which requires tumors to reach a detectable size, cfDNA detection from WGS data has been shown to provide improved sensitivity and lead time for recurrence detection [15,53]. In the CRC patient cohort, ctDNAmer detected disease recurrence earlier than radiological imaging, in some cases, as early as the first postoperative cfDNA sample. This may indicate that ctDNAmer could provide earlier recurrence detection compared to imaging. However, in the CRC cohort, imaging was performed less frequently than ctDNA testing. Therefore, further analyses and direct detection time comparison could be performed in the future to confirm the improvement in the median detection time. Early detection proposes the possibility for timely clinical actions, such as increased monitoring or earlier intervention, potentially improving patient outcomes.

When compared to the cfDNA allele frequency of clonal SNVs called from aligned primary tumor data, ctDNAmer achieves slightly improved detection accuracy. Although advanced modeling methods that apply deep learning-based approaches, such as MRD-Detect [15] and MRD-EDGE [53], may further enhance sensitivity, ctDNAmer offers a computationally efficient alternative. Unlike traditional WGS workflows that include read alignment, ctDNAmer relies on k-mer count and set operations, which require less time and computing resources. Further, by separating the signal from the two noise components during tumor fraction estimation, ctDNAmer provides an intuitive and transparent interpretation of the results, avoiding the “black box” nature of deep learning techniques. Nevertheless, to improve detection sensitivity, the tumor fraction estimation model used by ctDNAmer could be extended to a more advanced modeling approach that applies the sequence context information that the k-mers capture and can potentially learn which k-mers are most likely to be observed in the cfDNA. Using a larger set of normal controls to filter likely germline variants could potentially also improve the performance.

The main limitation of ctDNAmer is that it does not provide information about the genomic positions of the k-mers nor the variant types they represent. However, if needed for downstream analyses, this information could be retrieved by aligning k-mers assigned to the tumor component to the reference genome. Future studies should also investigate how sequencing depth and tumor mutational burden differences impact ctDNAmer’s performance.

In summary, we have demonstrated that reference-free ctDNA detection from WGS data is feasible using k-mers that capture tumor-specific somatic variation. ctDNAmer enables accurate tumor fraction estimation, achieves earlier recurrence detection compared to radiological imaging, and performs on par with methods based on aligned data while maintaining computational efficiency. These findings highlight the potential of the k-mer-based approach for ctDNA analysis and applications for measuring baseline tumor fraction, postoperative monitoring, early recurrence detection, and treatment response monitoring. By providing a scalable and efficient alternative to alignment-based methods, ctDNAmer could contribute to the broader clinical adoption of WGS-based liquid biopsy approaches.

## Methods

### 1. Patients, samples, and sequencing data

The study included 114 (90 FRFR and 24 FFPE tumor tissue biopsies) UICC stage III CRC patients treated at seven Danish hospitals between November 2002 and February 2019. All patients received standard of care treatment and recurrence surveillance, encompassing curative intent tumor resection, adjuvant chemotherapy (5-fluorouracil and oxaliplatin) at the clinician’s discretion, and CT scans at minimum at 12 and 36 months after tumor resection. The study was performed in accordance with the Declaration of Helsinki, and approved by the Danish National Committee on Health Research Ethics (case no. 2208092). Plasma samples from 30 healthy individuals included in the germline union were available at the Colorectal Cancer Research Biobank at Aarhus University Hospital and plasma samples from 15 healthy individuals applied for empirical noise estimation were collected through the blood bank at Aarhus University Hospital. The patients, the clinical data, and WGS data have previously been reported by Frydendahl et al. (2024) [54]. For details regarding sample processing and data generation, see Frydendahl et al. (2024) [54].

In brief, the tumor analysis was based on FRFR and FFPE tumor tissue biopsies with a minimum tumor fraction of 10 % (histological assessment). Patient blood samples were drawn before, after, and every third month for up to 3 years after the operation and processed to plasma within two hours of the blood draw. Cell-free DNA was extracted from 2 mL of plasma using the QiaAMP Circulating DNA kit (Qiagen). Normal DNA was extracted from peripheral blood mononuclear cells using QIAamp DNA Blood Kit (Qiagen). Tumor, Normal and plasma cell-free DNA samples underwent paired-end reads whole-genome sequencing (2 × 150 bp) using an Illumina NovaSeq platform. The target sequencing depths were 30X and 60X for tumors with tumor fractions above and below 30%, respectively, and 20X for cfDNA and normal DNA samples.

### 2. K-mer counting

ctDNAmer counts k-mers (k=51 by default; only odd-length k-mers used to avoid palindromes [55]) from the primary tumor, matched germline (DNA from peripheral blood mononuclear cells), and cfDNA sequencing reads using the KMC3 command-line tool [56]. Input data can be passed either in FASTQ or BAM format and all input files for each data source are processed simultaneously during counting. The KMC3 “cs” and “cx” parameters are set to 10^9^ and the “ci” to one, ensuring the inclusion of all k-mers in the output.

Before counting tumor k-mers we apply a quality filter by masking low-quality bases in tumor reads. Using the seqTK tool (version 1.4) or pysam (version 0.22.1), depending on the input format, nucleotide bases with a base quality score below 37 (maximum for Illumina NovaSeq) are replaced with “N”. Additionally, the last five bases from both ends of the reads, prone to higher error rates [57], are replaced with “N. Subsequently, KMC3 default exclusion of “N” bases ensures that regions with high error probability are not included in the tumor k-mer sets.

### 3. Identifying unique tumor k-mers

Unique tumor k-mer sets are created by subtracting the patient’s matching germline set from the tumor set. This step is followed by subtracting a union germline set from the tumor for additional filtering of germline sequences. ctDNAmer performs set subtraction with the KMC3 kmc_tools kmers_subtract command, which finds the difference between the two input sets based on k-mers, irrespective of k-mer counts. The resulting set is exported into a text file with the kmc_tools dump command. The text file represents the unique tumor k-mers (UT) set as a list of sequences in alphabetically sorted order and their respective counts in the tumor sequencing data.

For the CRC patient cohort analysis, we combined k-mers from 45 colorectal cancer patients’ germline k-mer sets and 30 healthy donor cfDNA k-mer sets to create a larger and more representative set of k-mers seen in the genomes of healthy cells. This germline union set combined across the 75 samples was used to identify unique tumor k-mers by subtracting the germline union set from the tumor set after matched germline k-mers have been removed (see Figure 1A dashed subpanel). Individual k-mer sets were combined into a single combined set, one set at a time, where in each step the union between two input sets was found using the KMC3 kmc_tools union operation.

#### 3.1. Filtering of the unique tumor k-mer sets

To increase the signal-to-noise ratio of the UT k-mer sets, ctDNAmer removes k-mers likely resulting from technical errors using two filters (Supplementary Figure 7). First, to avoid problems due to GC bias, characteristic of Illumina sequencing data, the GC content percentage of each UT k-mer is calculated, and all k-mers with a GC content below 20% or above 80% are removed. The GC content of each k-mer is calculated as the percentage of “G” and “C” bases in the k-mer by the following equation:

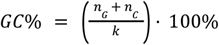

where *n*_*G*_ and *n*_*C*_ are the number of G and C bases in the k-mer, respectively, and *k* is the k-mer length.

Second, ctDNAmer filters k-mers based on their count in the tumor data. The aim of this is to remove k-mers seen only a few times or unusually many times in the tumor, as these k-mers are more likely to be created by sequencing errors. Furthermore, reducing the UT-set by applying patient-specific minimum and maximum counts also helps limit the computation time during TF estimation while ensuring the reliability of the results. We first set initial minimum and maximum cutoffs based on the mean and standard deviation of all tumor k-mers. The tumor k-mers count distribution is a positively skewed bimodal distribution with peaks at a count of one and a higher count that varies across samples. To avoid biased estimates, ctDNAmer calculates the mean and standard deviation after filtering out tumor k-mers with a count of one or above 50. The initial cutoffs are then defined as follows:

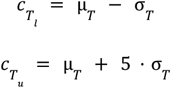

where 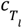 and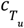 are the lower and upper cutoffs, respectively; μ_*T*_ is the mean count of tumor k-mers and σ_*T*_ is the standard deviation of tumor k-mer counts.

Next, ctDNAmer adjusts these initial cutoffs to ensure that each UT set exceeds the minimum UT set size threshold while remaining close to the target size. The minimum UT set size is set to 20,000 by default. This size threshold was determined based on a count-filtering test performed on a subset of the analyzed CRC patient cohort (see Methods section 3.2). To estimate TFs from WGS data with a different sequencing depth or for a different cancer type, the count-filtering test can be rerun as part of the ctDNAmer’s workflow to determine the corresponding optimal minimum UT set size.

If, after filtering based on the baseline count cutoffs, the resulting UT set exceeds the target size, the lower cutoff is incrementally increased to remove k-mers until further increase would reduce the set size below the required size or the lower cutoff reaches one count below the tumor mean. If the set size is still above the target size, the upper cutoff is incrementally decreased until further decrease would reduce the set size below the target size or until the upper cutoff reaches one count above the tumor mean. If the UT set size is still above the target after the cutoffs were adjusted to surround the tumor mean, both the lower and upper cutoffs are simultaneously increased by one until further increase would reduce the set size below the target.

Conversely, if the UT set contains fewer than the target number of k-mers after baseline filtering, the upper cutoff is first increased until the set size exceeds the target or the cutoff is equal to 100. If the upper cutoff reaches 100 and the set size is still below the target, the lower cutoff is decreased until the set size exceeds the target or the cutoff is equal to three. If the UT set size is still below the target, ctDNAmer retains the set below the target size, as k-mers observed only once or twice are unreliable due to the high probability of technical errors. An overview of the UT set filtering rules is provided in Supplementary Figure 7.

#### 3.2. Selecting the minimum UT k-mer set size

To assess the dependence of TF estimates on UT k-mer set size, the ctDNAmer tool can run an experiment testing different set sizes. The test is initialized with a baseline UT set that contains candidate UT k-mers with a count above five. We then test a grid of different set sizes by incrementally increasing the lower count cutoff. TF estimation is performed for at least one ctDNA-positive and one ctDNA-negative cfDNA sample from each patient using each count-filtered set.

We ran the count-filtering experiment on nine non-recurring patients from the CRC patient cohort. For each patient, TF was estimated for the preoperative (ctDNA-positive) and the last available postoperative (ctDNA-negative) cfDNA sample using each count-filtered set. Supplementary Figure 8 shows the TF estimates for the baseline UT set and all count-filtered sets. Based on the visual inspection of the TF estimates difference, 20,000 was chosen as the minimum UT set size for reliable TF estimation.

### 4. Annotation of unique tumor k-mer sets with cfDNA counts

ctDNAmer annotates the UT set with the cfDNA k-mer counts in two steps. First, the kmc_tools intersect command finds the intersection of the UT and the cfDNA set, annotating each UT k-mer with its corresponding count from the cfDNA set. Second, UT k-mers that are not observed in the cfDNA set are assigned a cfDNA count of zero.

### 5. Modeling the sample-specific background noise

To estimate the set-specific noise level of UT k-mers, ctDNAmer utilizes unmatched cfDNA samples. The k-mers are counted from the unmatched cfDNA samples in the same manner as the patients’ matched cfDNA k-mer sets. ctDNAmer then applies the kmc_tools intersect command to identify intersections between each unmatched cfDNA k-mer set and the UT set, and the count in the cfDNA set is recorded for each k-mer in the intersection. UT k-mers not included in the intersection are added to each set with a cfDNA count of zero. This ensures that k-mers unobserved in the unmatched cfDNA weigh the noise rate estimate towards zero. If the intersection is a null set, a set containing one pseudo-observation with a cfDNA count of one is created. Finally, the count data of all intersection sets are combined. In addition to the unmatched cfDNA count, each k-mer is tagged with the mean k-mer count of the specific unmatched cfDNA set in which it was observed.

The mean k-mer count is calculated based on all unmatched cfDNA k-mers after filtering out k-mers with low and high counts to avoid biasing the estimates. If the k-mers have a positively skewed bimodal distribution with a lower peak at a count of one, the lower cutoff is set as the count with the smallest number of k-mers between the two peaks. If the distribution is unimodal, with a single peak at a count of one, the lower cutoff is set to two. After applying the lower cutoff, the upper cutoff is selected based on the distribution of the remaining k-mers. ctDNAmer sets the upper cutoff as the smallest count above the second peak, if present, with less than 0.5% of the remaining k-mers.

The empirical set-specific noise distributions are modeled with a negative binomial distribution, where the mean noise estimate is scaled by the cfDNA set mean count to ensure correction for varying sequence depths. The model has the following form:

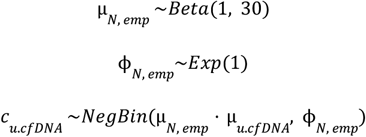

where μ_*N, emp*_ is the mean of the empirical noise rate, ϕ_*N, emp*_ is the variance scaling parameter (the inverse of parameter ϕ_*N, emp*_ scaled by the square of the mean μ_*N, emp*_^2^ controls the overdispersion, i.e. 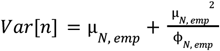[58]), *c*_*u*.*cfDNA*_ is the k-mer count in the unmatched cfDNA, and μ_*u*.*cfDNA*_ is the mean k-mer count of the respective unmatched cfDNA set.

The modeling is performed by setting the initial value of μ to 0.01 and the initial value of ϕ_*N, emp*_ to 0.1 for all chains. After 100 burn-in iterations, 1000 samples from the posterior distributions of μ_*N, emp*_ and ϕ_*N, emp*_ are obtained from four chains, and mean values of the posterior samples are saved as the parameters’ estimates. The model convergence assessment is done based on the same statistics and visualizations as for the tumor fraction estimation (see Methods section 6).

### 6. Tumor fraction estimation

ctDNAmer estimates the circulating tumor fraction of the cfDNA sample by probabilistic modeling of the UT k-mers’ cfDNA counts. The TF is expected to affect the k-mer count only if the k-mer contains a tumor-specific somatic variant, not if the k-mer is a missed germline k-mer or represents background noise. Therefore, cfDNA counts are modeled with a three-component mixture model (ctDNA, germline, noise), and each k-mer is assigned to one of the three components based on the component weights. All mixture components are modeled with negative binomial distributions.

The means of the mixture component distributions are set based on prior expectations. The ctDNA component mean is set to *TF* · μ_*cfDNA, GC*_ as it is expected to depend on the TF. Here, the cfDNA mean, μ_*cfDNA, GC*_, is the mean count of k-mers with GC-content GC in the cfDNA. Similarly, the germline component mean is set to 0. 5 · μ _*cfDNA, GC*_, as the majority of missed germline k-mers are expected to represent heterozygous germline variants with allele frequency close to 0.5. The noise component mean is set to μ_*N, emp*_ · μ_*cfDNA, GC*_, where μ_*N, emp*_ is the noise rate estimated from empirical noise data (see Methods section 5). The variances of the mixture components are based on the distribution means and variance scaling parameters: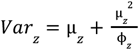[58], where μ_*z*_ is the distribution mean and ϕ_*z*_ is the variance scaling parameter.

All three components’ mean count estimates are scaled by the mean k-mer count of the cfDNA sample, μ_*cfDNA, GC*_, to ensure correction for varying sequence depths of the cfDNA samples. As the average sequencing depths are unknown because sequencing reads are not aligned to the reference genome, the mean cfDNA count of matched germline k-mers, which highly correlates with the mean sequencing depth of the cfDNA samples (see the correlation for the analyzed CRC cohort in Supplementary Figure 9a), is used for component mean estimate adjustment as a proxy for sequencing depth. The μ_*cfDNA, GC*_ estimates used during modeling are calculated separately for each GC content value to account for the coverage GC bias inherent in Illumina data.

The mean cfDNA count is calculated based on counts of cfDNA k-mers observed in the patient’s germline set to avoid bias from ctDNA k-mers and technical noise. First, the kmc_tools intersect command finds the intersection between the cfDNA and the germline set. After calculating the GC content of the intersection k-mers, the count distribution of each subset of k-mers is constructed by counting k-mers between a set-specific minimum cutoff and 100. K-mers with counts below the minimum cutoff or above 100 are excluded to minimize bias from technical noise, amplified genomic regions, or k-mers that are not uniquely positioned in the genome. The minimum count cutoff is determined similarly to the unmatched cfDNA k-mers minimum cutoff (see Methods section 5). Briefly, it is set as the count with the smallest number of k-mers between the two peaks of the distribution for a positively skewed bimodal count distribution, or to two for a uniformly decreasing count distribution.

The probabilistic model fitted to the UT k-mers cfDNA count data has the following form:

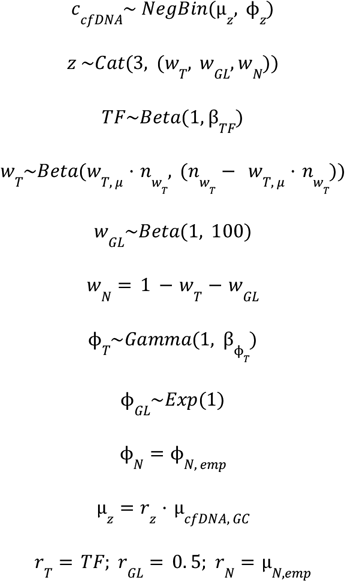

where *c*_*cfDNA*_ is the k-mer count in the cfDNA, μ_*z*_ and ϕ_*z*_ are negative binomial distribution parameters as described above, *r* indicates the component mean scaling factor as described above, *z* ∈ {*T, GL, N*} indicates the component (T: ctDNA; GL: germline; N: noise), *w*_*T*_, *w* _*GL*_ and *w*_*N*_ are the ctDNA, germline, and noise component weights, respectively, *TF* is the tumor fraction of the cfDNA sample, β_*TF*_ is the beta parameter of the *TF* prior Beta distribution, *w*_*T, μ*_ is the mean of the tumor component weight prior distribution,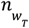is the sample size of the tumor component weight prior distribution,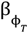is the ϕ_*T*_ parameter prior Gamma distribution beta parameter, μ_*N, emp*_ and ϕ_*N, emp*_ are the noise distribution mean and variance scaling parameter estimated from empirical data, and μ_*cfDNA, GC*_ is the mean count of cfDNA k-mers with matching GC content.

For the ϕ _*GL*_ and ϕ_*T*_ parameters, lower bounds are set based on upper bounds of the respective component variances. The maximum variance of the germline component is calculated based on the cfDNA sample variance,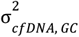, estimated based on matched germline k-mers’ cfDNA counts. The cfDNA k-mers’ variance strongly correlated with the sequencing depth variance in the analysed CRC patient cohort (Supplementary Figure 9b). The lower bound for the germline component variance scaling parameter ϕ_*GL*_ is set to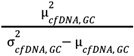, ensuring that the component variance will not exceed the variance of the cfDNA k-mers observed in the matched germline set. The GC content value used for the lower bound estimation is chosen as *GC* = *argmax*_20<*GC*<80_ (μ_*cfDNA, GC*_). The tumor component variance scaling parameter ϕ is constrained with a lower bound that sets the maximum variance to be larger than the component mean based on the scaling factor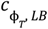 as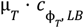. This ensures that no convergence issues arise due to equally good model fits resulting from a high TF estimate with a low tumor component variance and a low TF estimate with a high tumor component variance.

Similarly to empirical Bayes methods, where prior distribution parameters are estimated from the data, ctDNAmer chooses between two sets of prior distribution parameters based on the cfDNA mean count of the UT k-mers that are observed in the cfDNA set. If the mean count is below the germline component distribution mean 0. 5 · μ_*cfDNA, GC*_, there is no clear evidence of ctDNA presence, and prior distribution parameters are set to be indicative of low TF 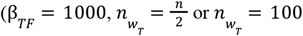, for baseline or subsequent cfDNA samples, respectively;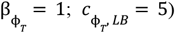. Conversely, UT k-mers’ mean above 0. 5 · μ_*cfDNA, GC*_ can indicate high TF, and prior distribution parameters are set accordingly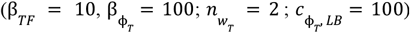. If convergence issues occur, the 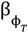parameter is increased to 1000.

The tumor component weight prior distribution mean *w*_*T, μ*_ is set to 50% in the baseline cfDNA samples and to the baseline sample mean *w*_*T*_ estimate in all subsequent samples. Also, a lower bound of 1% is set for *w*_*T*_ to ensure that a small fraction of the k-mers is always assigned to the tumor component. This prevents TF estimation based on a small number of k-mers, reducing the associated uncertainty. For example, given the minimum UT set size of 20,000, the lower bound ensures the assignment of approximately 200 k-mers to the tumor component.

For the baseline cfDNA samples, the germline component weight *w*_*GL*_ is estimated using a prior Beta distribution with parameters 1 and 100. As the number of germline k-mers mistakenly included in the UT sets is not expected to change, the mean germline weight estimate *w*_*GL*_ and the variance scaling parameter ϕ_*GL*_ calculated from the baseline cfDNA sample are applied as fixed parameters for the TF estimation of all subsequent samples.

The probabilistic models are implemented in STAN version 2.35 [58]. Using the rstan package (version 2.32.6) with the seed set to one, 2000 posterior samples are obtained from four chains, after 500 burn-in iterations. The rstan summary function is used to obtain the mean, 95% credible intervals, Gelman-Rubin convergence diagnostic statistic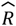, and the effective sample size of the *TF, w*_*T*_, *w*_*GL*_, ϕ_*T*_ and ϕ_*GL*_ parameters based on the total 8000 posterior samples.

We assessed model convergence for the CRC patient cohort data by visually inspecting the trace plots to confirm that there were no long-range trends and that the samples were most likely drawn from the posterior distributions. In addition, the Gelman-Rubin convergence diagnostic statistic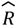was used to assess that all chains had converged. The autocorrelation plot, generated with the rstan function stan_ac, and the effective sample size were used to confirm minimal autocorrelation between the posterior samples.

### 7. Estimating the ctDNA status of the cfDNA samples

We set the detection cutoff by maximizing Youden’s J statistic [65], indicated by the dashed blue line in Figure 2a. The true ctDNA status of the samples was identified from the standard-of-care radiological follow-up results. All preoperative samples were classified as ctDNA-positive. In addition, postoperative samples of recurring patients (recurrence confirmed by radiological imaging) were classified as ctDNA-positive if collected after the end of initial treatment (surgery or adjuvant chemotherapy) and before the start of the last recurrence treatment, if recurrence treatment was administered. Postoperative samples of non-recurring patients (no abnormalities detected on imaging) collected after the last treatment, either surgery or adjuvant chemotherapy, were classified as ctDNA-negative. This classification resulted in 207 positive and 511 negative ctDNA samples.

If the posterior mean of the TF was equal to or exceeded the predefined detection cutoff, the cfDNA sample was classified as ctDNA-positive, and the mean TF estimate was reported. Conversely, if the TF was below the cutoff, the cfDNA sample was classified as ctDNA-negative, and the TF estimate was set to zero.

### 8. Detection of clonal SNVs in cfDNA

#### 8.1. Identifying clonal SNVs and quality filtering of the variant set

We used the cfDNA allele frequency of clonal SNVs, identified from variant sets called from the aligned sequencing data, as an independent TF measure for comparison with the ctDNAmer method.

First, sequencing adapters were trimmed with cutadapt, and reads were aligned to the hg38 reference genome using bwa-mem. Duplicates were marked with GATK’s MarkDuplicates command and variants were called using Mutect2 tumor-normal mode [59]. Subsequently, variant sets were filtered using the FilterMutectCalls command with the following settings: max-events-in-region set to three, min-slippage-length set to eight, and normal-p-value-threshold set to 0.0001. We further filtered Mutect2 calls with vcftools (version 0.1.16) to retain only SNV variants (indels removed) that had the ‘PASS’ FILTER flag and were located on autosomes or allosomes. The variants were then annotated using Ensembl Variant Effect Predictor (VEP, version 105.0) [60]. From the VEP output, we extracted the coverage at variant positions, the number of reads with the alternative allele, variant impact, and gene labels. To create the somatic mutations input data set for CNAqc, we tagged variants in coding regions with a ‘high’ impact annotation as potential drivers, recorded their gene labels, and calculated variant allele frequency by dividing the number of reads with the alternative allele by the position coverage.

To create the copy number segment data for CNAqc, we used the sequenza toolset (sequenza R package and Sequenza-utils version 3.0.0) [61]. First, a GC wiggle track was created from the reference genome with a window size of 50. Next, a seqz file was generated for each tumor sample using the bam2seqz command with the qformat flag set to “illumina”. The seqz file was then binned with the seqz_binning command with a window size of 50. The binned file was processed using the sequenza R package sequenza.extract function (gamma = 280, kmin = 200, max.mut.types = 1, min.reads.baf = 5, and min.reads = 10). Data was extracted from autosomes and chromosome X for female patients, and only from autosomes for male patients. Next, a grid search of the parameter space was performed using the sequenza.fit function. Cellularity (tumor purity) and ploidy candidate values were set based on predefined cutoffs: cellularity values were chosen from the range of 0.1 to 0.85 with a step size of 0.01, and ploidy values from the range of 1 to 2.5 with a step size of 0.1. These cutoffs were adjusted per sample if the CNAqc peaks analysis failed.

To perform quality control on the variant data, we ran CNAqc [62] (version 1.0.0) on the annotated SNV calls, copy number segments, and tumor purity and ploidy estimates obtained from sequenza. We used the CNAqc analyze_peaks function, which compares peaks observed in variant allele frequency data to theoretical expectations. The data quality was confirmed if the peaks’ analysis was passed for diploid heterozygous (1:1 genotype) variants and other simple copy number states with a considerable number of variants (> 5%). SNVs in diploid heterozygous copy number states (clonal SNVs) that passed quality control were selected for downstream analyses.

We further filtered the clonal SNV set by excluding variants with frequencies below and above specified cutoffs. The cutoff for low-frequency variants was found by analyzing the allele frequency distribution. By default, if the distribution was bimodal, the cutoff was set at the lower frequency peak; otherwise, it was set to 0.1. The lower cutoff was further adjusted if peak detection or clonal variant selection failed. The upper cutoff was set to 0.9 by default and adjusted to a lower sample-specific cutoff to exclude outliers if clonal variant selection failed. Next, we used the MOBSTER R package (version 1.0.0) [63] to separate clonal variants from noise and passenger variants at lower frequencies. The mobster_fit function with default settings was applied to find the best-fitting model for the variant data. Cluster assignments were extracted using the Clusters function, with a cutoff_assignment set to 0.85 and adjusted to 0.5 if no variants passed the default cutoff. Variants assigned to the C1 cluster, which is characterized by the highest (Beta) mean and in diploid regions represent clonal mutations, were selected for tracking in cfDNA.

Before searching for clonal SNVs in cfDNA, we filtered variants based on the primary tumor reads alignment information to exclude variants from low-quality regions and positions. Using pysam (version 0.22.1), a pileup was constructed at each variant position, and all variants that did not pass a set of quality requirements were removed. Specifically, only one alternative allele matching the alternative called by Mutect2 was allowed at each position. At least 90% of aligned reads needed to be high-quality, with a minimum of 20 high-quality reads in total and at least two high-quality reads with the alternative allele. In addition, a median base quality above 20, and a median distance from the variant position to the read end greater than five were required across the aligned reads. A read was defined as high-quality if it was not a secondary or a supplementary alignment, had no indels or clippings, contained fewer than two mismatches with the reference genome across the read (allowing only the target variant as a mismatch), and had a mapping quality of 60. Clonal SNVs passing these filters were saved for ctDNA detection (median of 1387 clonal SNVs per patient; range 126 - 52,475 SNVs).

#### 8.2. Tracking clonal SNVs in cfDNA

To track clonal SNVs in cfDNA, we used pysam to create a pileup at each variant position and assess a set of quality requirements. As with the primary tumor data, we required that only the alternative called by Mutect2 can be observed, with no other alternative alleles present. In addition, 90% of the aligned reads had to be of high quality, with at least 15 high-quality reads covering the position. The definition of a high-quality read was unchanged from the primary tumor data analysis. We also required a median base quality of 20 and a median end distance of at least five at the position, similar to the primary tumor quality requirements. If all quality filters were passed, the variant and its allele frequency information were saved. Finally, we calculated the mean allele frequency of the variants and compared it with the TF estimate obtained with ctDNAmer.

### 9. Tumor fraction estimation with ddPCR and Signatera

Data on ctDNA detection were available from previous studies using ddPCR [50] and Signatera [51], respectively. The full methodology is described in the respective papers. In brief, cfDNA was extracted from 8mL of plasma for both methods and whole-exome sequencing was conducted on the primary tumor to inform target selection for ctDNA analysis. For ddPCR, a single clonal variant was selected per patient. Analysis was conducted using a duplex assaying targeting the mutation and corresponding wildtype to calculate the ctDNA allele frequency. For Signatera, 16 clonal variants were targeted by multiplex PCR for each patient. Each variant was quantified in the plasma through ultradeep targeted sequencing, and the mean allele frequency of the 16 targets were used to calculate the overall tumor fraction.

### 10. Statistical analysis and code availability

Statistical analysis was performed with Python 3.12 and R version 4.4.1. All the workflows were built and run using the Snakemake workflow management tool version 8.25.2 [64]. Workflows and analytic code used for this work are available at https://github.com/BesenbacherLab/ctDNAmer.

## Data availability

The clinical and WGS data used in the study are available through controlled access from GenomeDK (https://genome.au.dk/library/GDK000005/). To protect the privacy and confidentiality of patients in this study, personal data, including clinical and sequence data, are not made publicly available in a repository or the supplementary material of the article. The data can be requested at any time from the corresponding authors. Any requests will be reviewed within a time frame of 2 to 3 weeks by the data assessment committee to verify whether the request is subject to any intellectual property or confidentiality obligations. All data shared will be de-identified. Request for access to raw sequencing data furthermore requires that the purpose of the data re-analysis is approved by The Danish National Committee on Health Research Ethics.

## Declaration of competing interest

The authors declare the following financial interests/personal relationships, which may be considered as potential competing interests. CLA reports sponsored research agreements with Natera, Bio-Rad Laboratories, AccuraGen Inc., and Labcorp.

## Acknowledgments

Patient recruitment and all the clinical info collection was done by the IMRPOVE-consortia. Following is the IMPROVE-consortia (listed alphabetically): Alessio Monti (Department of Surgery, North Denmark Regional Hospital Hjørring, Hjørring, Denmark), Claudia Jaensch (Department of Surgery, Regional Hospital Gødstrup, Herning, Denmark), Ismail Gögenur (Center for Surgical Sciences, Zealand University Hospital, Køge, Denmark), Jeppe Kildsig (Department of Surgery, Copenhagen University Hospital, Herlev, Denmark), Kåre Andersson Gotschalck (Department of Surgery, Regional Hospital Horsens, Horsens, Denmark), Lene Hjerrild Iversen (Department of Surgery, Aarhus University Hospital, Aarhus, Denmark), Nis Hallundbæk Schlesinger (Department of Surgery, Copenhagen University Hospital, Bispebjerg, Denmark), Ole Thorlacius-Ussing (Clinical Cancer Research Center, Aalborg University, Aalborg, Denmark), Per Vadgaard Andersen (Department of Surgery, Odense University Hospital, Odense, Denmark), Peter Bondeven (Department of Surgery, Regional Hospital Randers, Randers, Denmark),Thomas Kolbro (Department of Surgery, Odense University Hospital, Svendborg, Denmark), Uffe Schou Løve (Department of Surgery, Regional Hospital Viborg, Viborg, Denmark).

We extend our thanks to the patients and their families. This study was supported by the Novo Nordisk Foundation [grant numbers NNF21OC0069056 (SB), NNF21OC0069105 (SB), NNF17OC0025052 (CLA) and NNF22OC0074415 (CLA)] and the Danish Cancer Society [grant number R257-A14700 (CLA)]. The opinions, results, and conclusions reported in this article are those of the authors and are independent of funding.

## Supplementary material

**Supplementary Figure 1.**
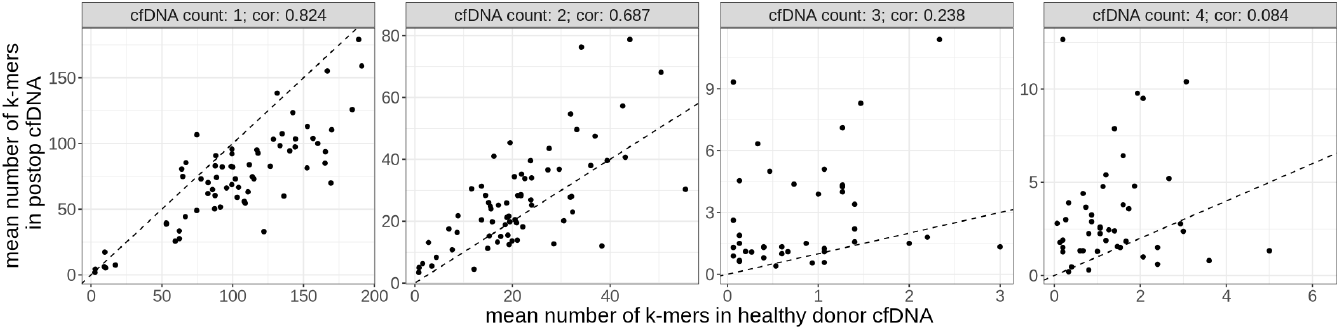
The mean number of UT k-mers in 15 healthy donor cfDNA samples compared with the mean number of UT k-mers in the ctDNA-negative postoperative cfDNA samples of the respective patients. The analysis is stratified based on the k-mer count in the two types of cfDNA samples. Only non-recurring patients and postoperative samples obtained after the end of the final treatment (either surgery or adjuvant chemotherapy) were included in the analysis.

**Supplementary Figure 2.**
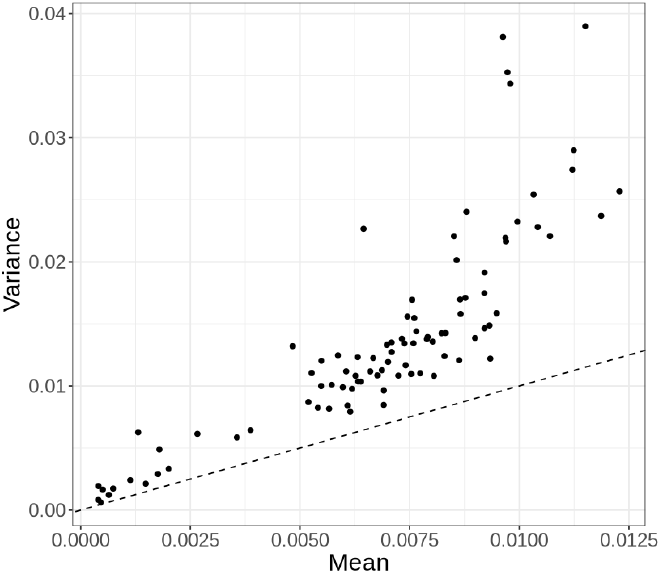
The mean and variance of the estimated background noise distributions. The distribution means are scaled by assuming a fixed cfDNA mean count of 30. The dashed black line indicates a Poisson distribution with mean equal to variance.

**Supplementary Figure 3.**
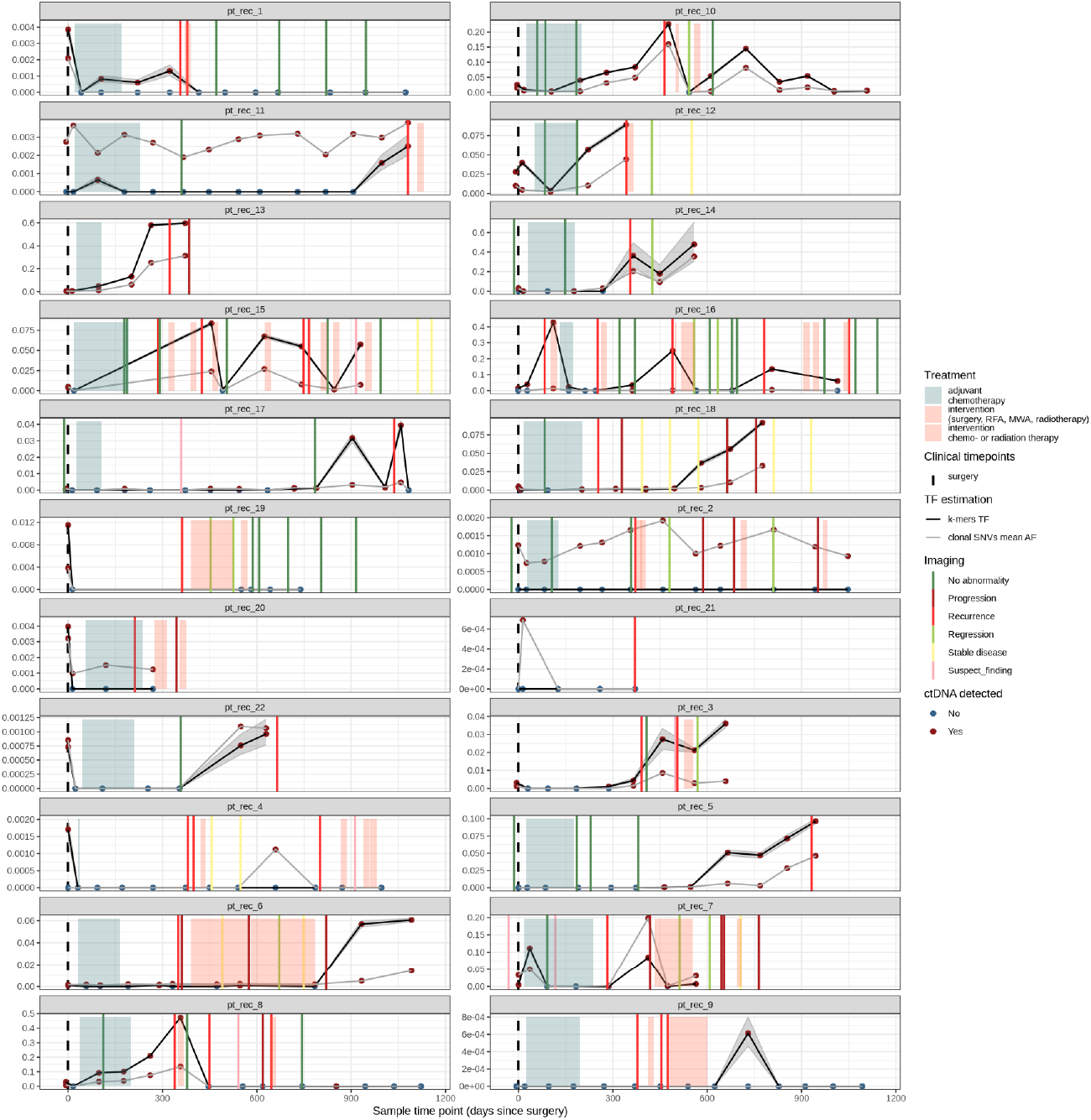
TF estimates of the recurring patients and comparison to clonal SNVs mean allele frequencies. Each panel represents a patient timeline. The day of the surgery (dashed black line) is marked as zero on the x-axis. Y-axis limits vary across subpanels.

**Supplementary Figure 4.**
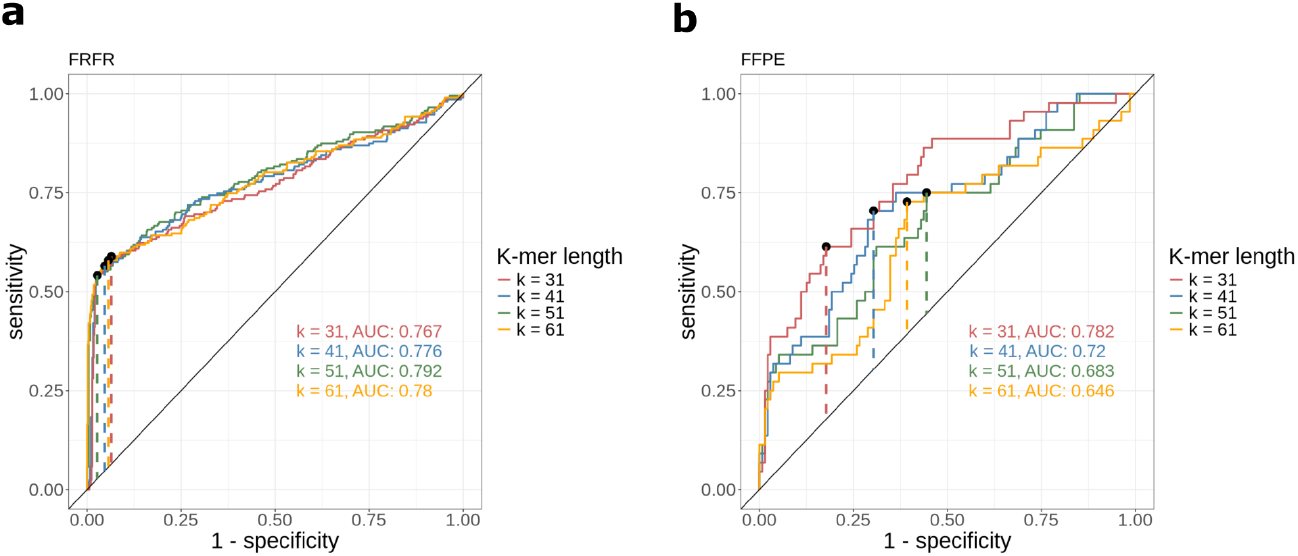
Comparison of ctDNA detection results across different k-mer lengths. Dashed lines indicate the maximum Youden J’s, used to determine ctDNA detection cutoff. **a)** ROC curves of the 90 patients with FRFR tumor tissue samples; **b)** ROC curves of the 24 patients with FFPE tumor tissue samples.

**Supplementary Figure 5.**
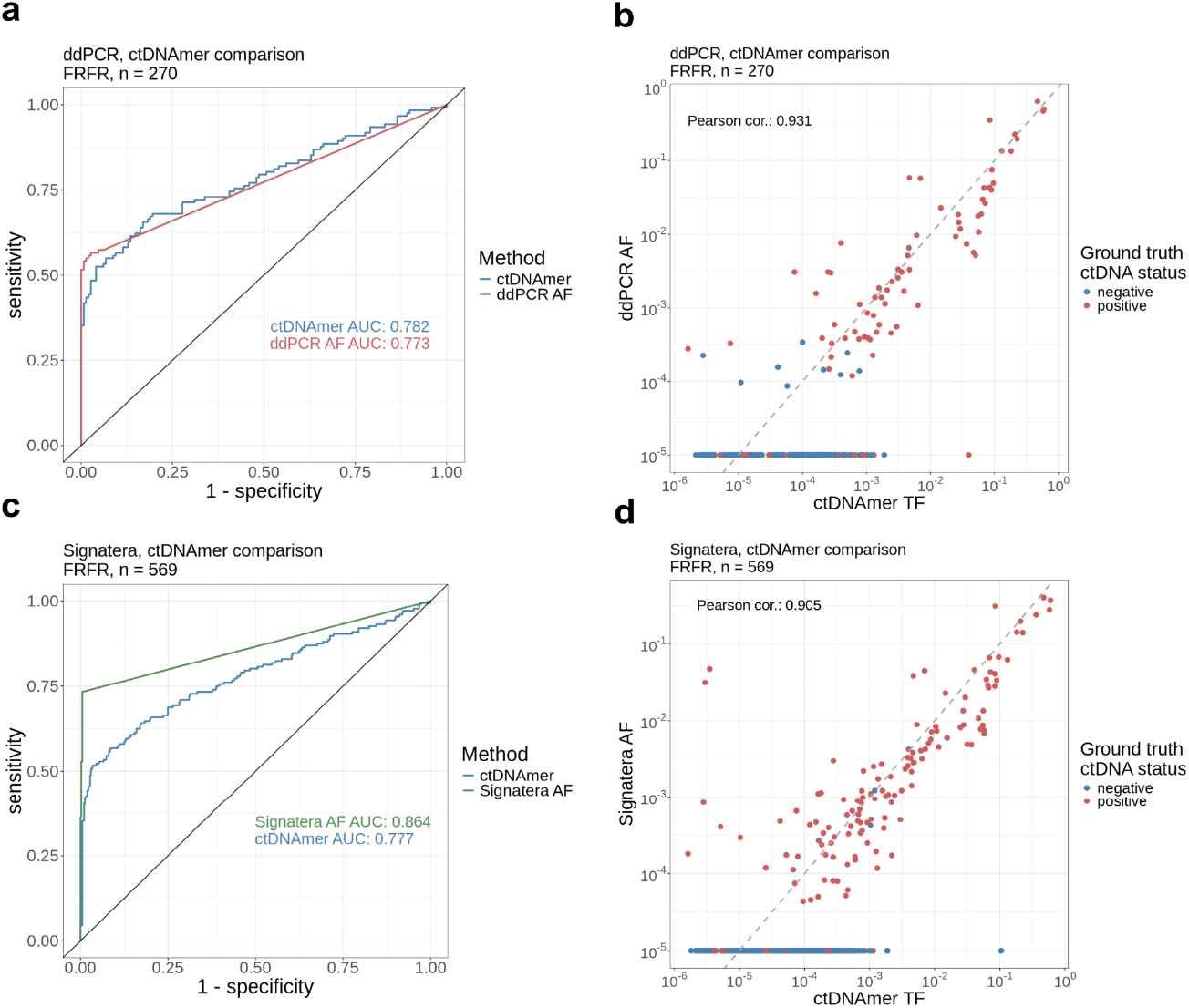
Comparison of ctDNAmer with ddPCR and Signatera methods. Only samples with known ground truth status and estimates available from both methods were included in the analysis. The number of included cfDNA samples is shown in each subplot title. **a)** ROC curve comparing ctDNA detection based on FRFR tumor samples with ctDNAmer and ddPCR; **b)** Comparison of ctDNAmer’s TF estimates and ddPCR allele frequencies, axes are on a logarithmic scale, ddPCR AF zero values are changed to 10^−5^; **c)** ROC curve comparing ctDNA detection based on FRFR tumor samples with ctDNAmer and Signatera; **d)** Comparison of ctDNAmer’s TF estimates and Signatera allele frequencies, axes are on a logarithmic scale, Signatera AF zero values are changed to 10^−5^. AF: allele frequency.

**Supplementary Figure 6.**
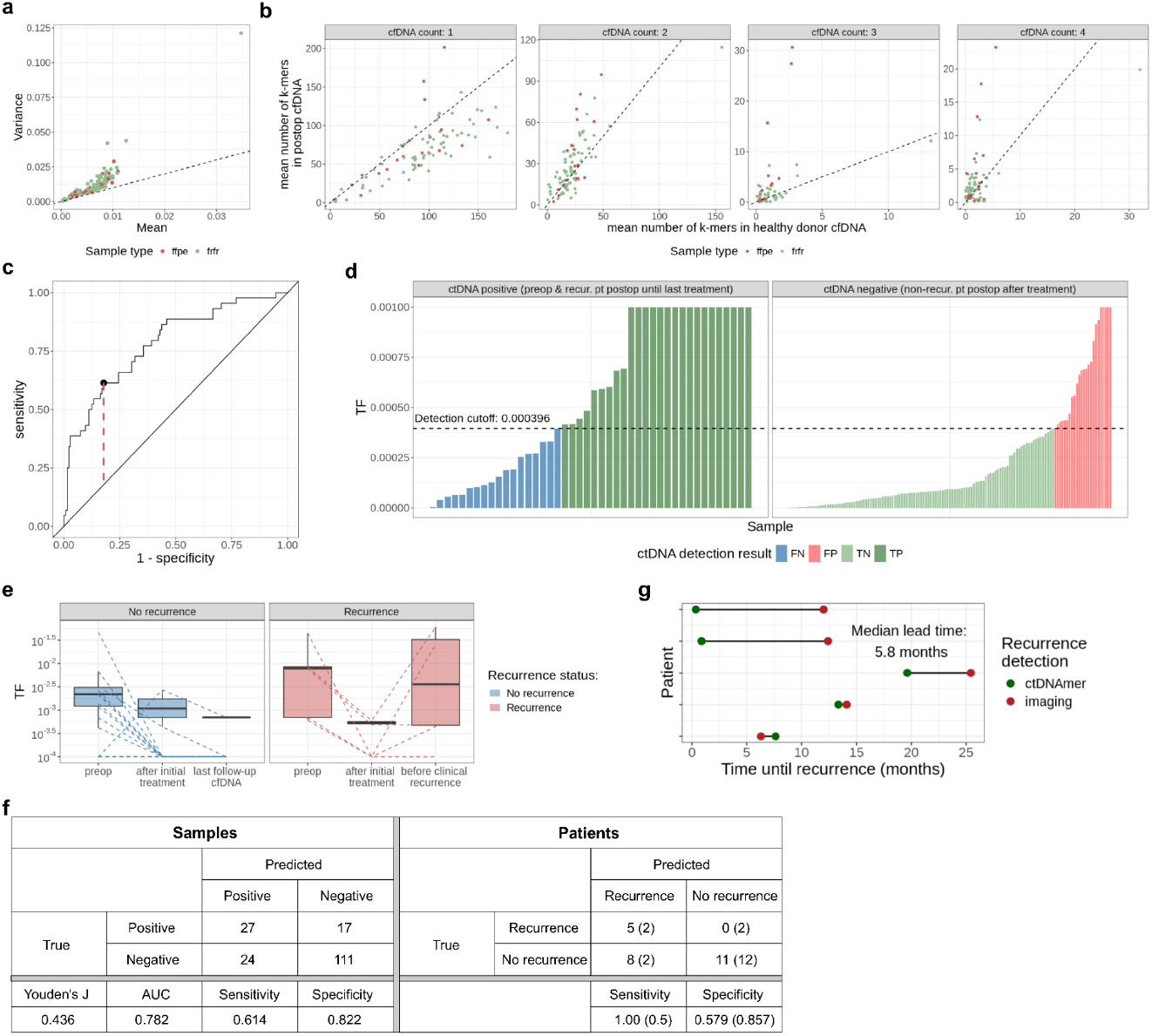
Results on the FFPE cohort (n=24) with k = 31. **a)** Background noise distribution parameter in comparison with FRFR cohort. The distribution means are scaled by assuming a fixed cfDNA mean count of 30. The dashed black line indicates a Poisson distribution with mean equal to variance; **b)** Mean number of UT k-mers in healthy cfDNA compared to the mean number of k-mers in ctDNA-negative postoperative samples of non-recurring patients in comparison with FRFR cohort; **c)** ROC curve. The dashed red line indicates the maximum Youden’s J statistic value; **d)** Estimated tumor fractions of ground truth ctDNA-positive and -negative samples. Samples are colored based on their detection result. The dashed black line indicates the detection cutoff chosen based on maximizing Youden’s J. FN: False negative; FP: False positive; TN: True negative; TP: True positive; **e)** Comparison of TF estimates of recurring and non-recurring patients at different categorical time points; dashed lines indicate TF estimates of each individual across the time points, y-axis is on a logarithmic scale, zero values were excluded from the boxplot calculation and set to 10^−4^ for the dashed lines; **f)** Confusion matrices of detection results on the sample and patient level. The values outside the parentheses in the patient matrix represent serial analysis that includes all samples collected during the follow-up period. The values inside the parentheses in the patient matrix represent landmark analysis where only one sample, the first sample collected after the surgery and before the start of the adjuvant chemotherapy, was analysed from each patient (18 out of the 24 patients had the landmark sample available); **g)** Comparison of recurrence detection times between ctDNAmer and radiological imaging. The ctDNAmer’s recurrence detection time point was chosen as the first postoperative ctDNA sample where ctDNA was detected, even if adjuvant chemotherapy was still ongoing.

**Supplementary Figure 7.**
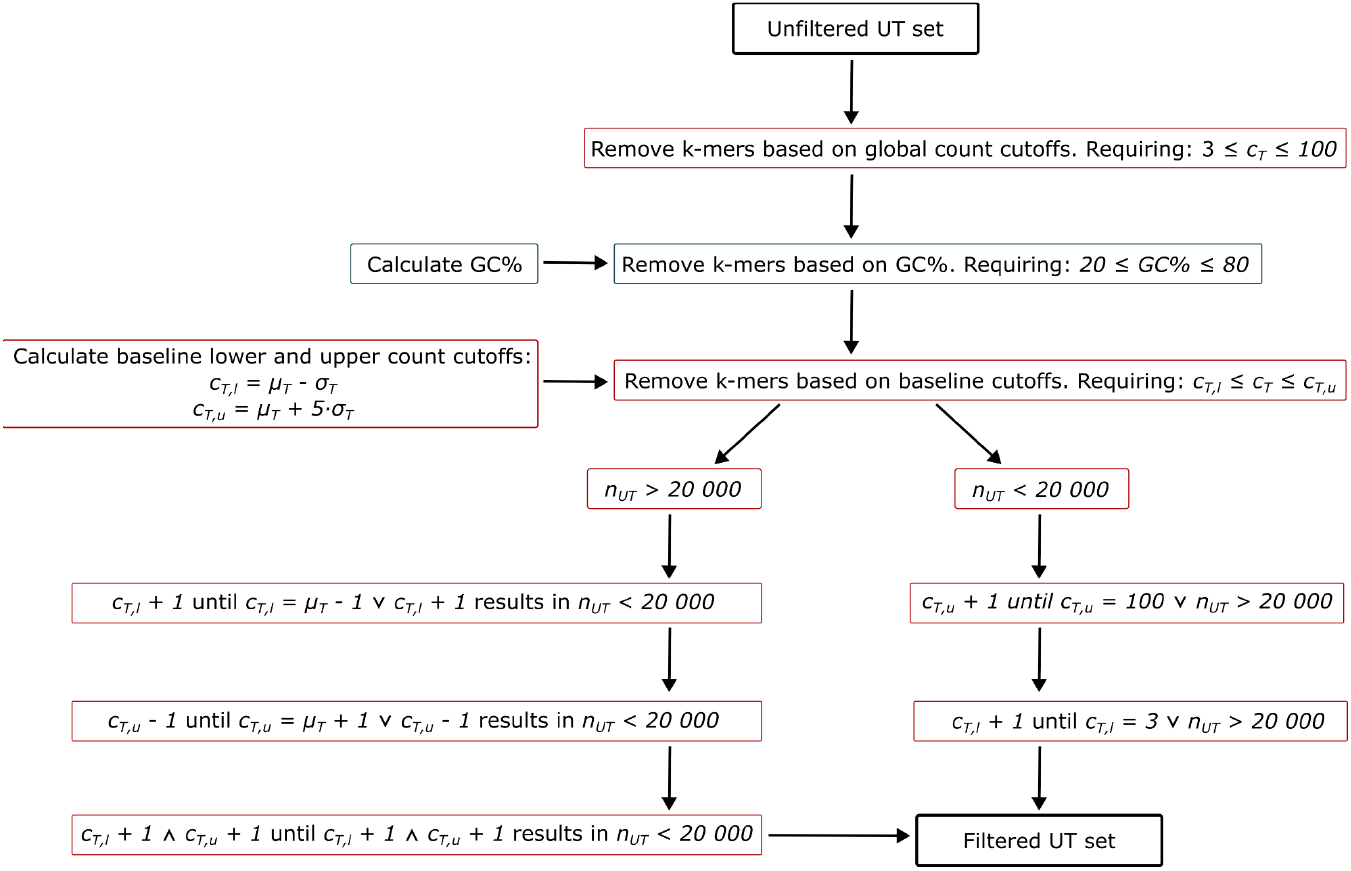
Schema for filtering the unique tumor sets based on the GC content (boxes with blue border) and tumor count (boxes with red border). UT: unique tumor, c_T_: tumor k-mer count, c_T,l_: tumor k-mer count lower cutoff, c_T,u_: tumor k-mer count upper cutoff, μ_T_: tumor k-mers count distribution mean, σ_T_: tumor k-mers count distribution standard deviation, n_UT_: number of unique tumor k-mers.

**Supplementary Figure 8.**
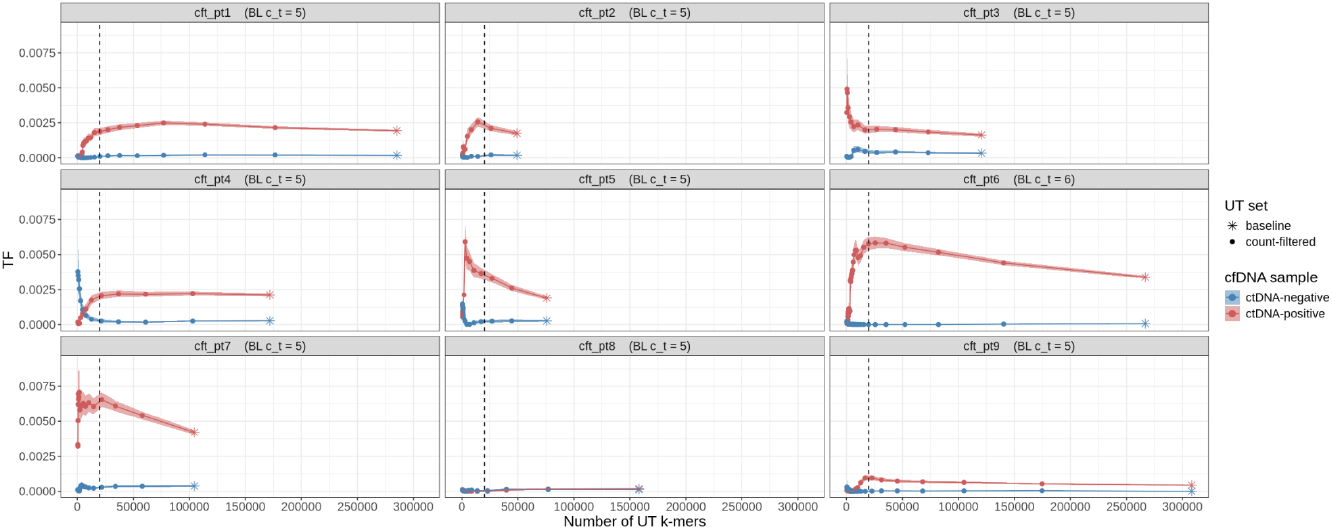
Tumor fraction estimates for nine non-recurring patients across UT sets of different sizes. UT set sizes vary across patients as filtering was done based on increasing the lower count cutoff and identical cutoffs resulted in UT sets of different sizes. The data points indicate mean tumor fraction estimates and colored ribbons around the TF mean estimates show 95% credible intervals. The black vertical dashed lines indicate the chosen minimum UT set size of 20,000. BL: baseline; c_t: minimum tumor count in the largest tested (baseline) set.

**Supplementary Figure 9.**
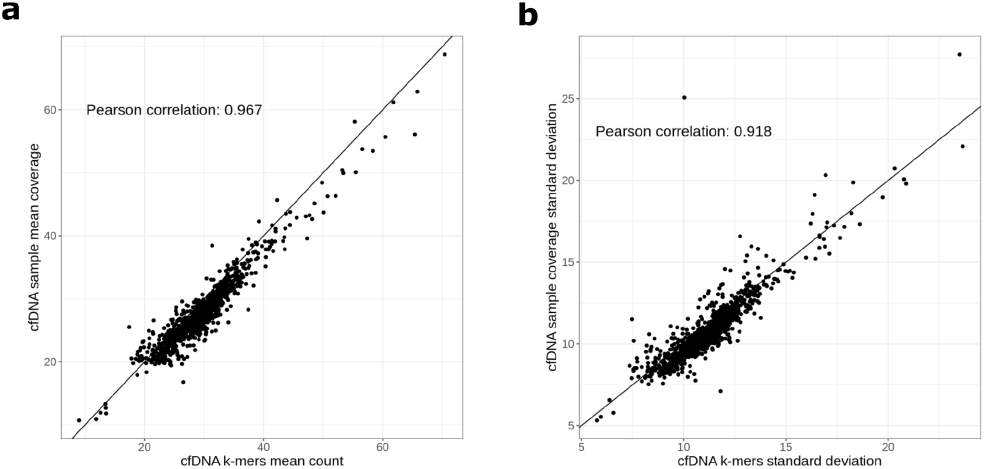
**a)** cfDNA sample mean coverage compared to the mean cfDNA count of matched germline k-mers; **b)** standard deviation of cfDNA sample coverage compared to the standard deviation of the matched germline k-mers’ cfDNA count.

